# A pooled sequencing approach identifies a candidate meiotic driver in Drosophila

**DOI:** 10.1101/106336

**Authors:** Kevin H-C Wei, Hemakumar M. Reddy, Chandramouli Rathnam, Jimin Lee, Deanna Lin, Shuqing Ji, James M. Mason, Andrew G. Clark, Daniel A. Barbash

**Author notes:** Correspondence to: Dr. Daniel Barbash, 401 Biotechnology Bldg., Cornell Univ., Ithaca, NY 14853, Em.

## Abstract

Meiotic drive occurs when a selfish element increases its transmission frequency above the Mendelian ratio by hijacking the asymmetric divisions of female meiosis. Meiotic drive causes genomic conflict and potentially has a major impact on genome evolution, but only a few drive loci of large effect have been described. New methods to reliably detect meiotic drive are therefore needed, particularly for discovering moderate-strength drivers that are likely to be more prevalent in natural populations than strong drivers. Here we report an efficient method that uses sequencing of large pools of backcross (BC1) progeny to test for deviations from Mendelian segregation genome-wide of single-nucleotide polymorphisms (SNPs) that distinguish the parental strains. We show that meiotic drive can be detected by a characteristic pattern of decay in distortion of SNP frequencies, caused by recombination unlinking the driver from distal loci. We further show that control crosses allow allele-frequency distortion caused by meiotic drive to be distinguished from distortion resulting from developmental effects. We used this approach to test whether chromosomes with extreme telomere-length differences segregate at Mendelian ratios, as telomeric regions are a potential hotspot for meiotic drive due to their roles in meiotic segregation and multiple observations of high rates of telomere sequence evolution. Using four different pairings of long and short telomere strains, we find no evidence that extreme telomere-length variation causes meiotic drive in Drosophila. However, we identify one candidate meiotic driver in a centromere-linked region that shows an ~8% increase in transmission frequency, corresponding to a ~54:46 segregation ratio. Our results show that candidate meiotic drivers of moderate strength can be readily detected and localized in pools of F1 progeny.

## INTRODUCTION

Mendel’s Law of equal segregation is not without its exceptions. Large deviations from the expected 50:50 segregation ratio of alleles from heterozygous parents are not common, but they have been observed in many different organisms (Pardo-Manuel de Villena and Sapienza 2001; Lindholm *et al.* 2016). These deviations, referred to as meiotic drive and segregation distortion for those involving meiotic or post-meiotic processes, respectively, are of great interest for the same reasons one studies large-effect mutations: they reveal fundamental information about biological processes such as meiosis and gametogenesis through mechanisms that disrupt them. Non-Mendelian transmission also provides a window into evolutionary conflicts between host fitness and selfish alleles causing meiotic drive and segregation distortion. These conflicts can have significant impacts on genome evolution (Sandler and Novitski 1957) and have been implicated in speciation events (McDermott and Noor 2010).

The prevalence of alleles causing non-Mendelian segregation (henceforth referred to as distorters) in natural populations is unknown. One reason is that distorters are expected to rapidly reach fixation, and once fixed their ability to distort becomes impossible to detect as competing alleles have been driven to extinction. By extension, strong distorters are less likely to be segregating at intermediate frequencies than weak distorters given equal rates of emergence, because the speed of fixation is dependent on the strength of distortion (Thomson and Feldman 1976). However, strong distorters are easier to identify as they can be detected from moderate numbers of progeny. In fact, the small number of characterized distorters dramatically influence progeny ratios. For example, the *D* locus of *Mimulus guttatus* causes nearly 100% self-transmission when paired with a chromosome of a sister species lacking it (Fishman and Willis 2005). Another example is the *Segregation Distorter* gene (*SD*) in *Drosophila melanogaster*, where *SD*/+ heterozygous flies almost exclusively sire *SD* offspring (Larracuente and Presgraves 2012). However, despite their very strong distortion phenotypes, both *D* and *SD* are still segregating at intermediate population frequencies in natural populations because they have deleterious pleiotropic effects on fitness that counterbalance their rates of fixation (Wu *et al.* 1989; Fishman and Saunders 2008; Brand *et al.* 2015).

The paucity of known weak distorters runs counter to the expectation that they segregate in populations longer than strong distorters. This is likely due to detection bias, as one must screen very large numbers of progeny to overcome sampling effects for weak distorters. A further challenge is that small differential viability effects among progeny can either mask or falsely mimic non-Mendelian segregation. However, weak distorters are of great interest as they may represent the initial evolutionary stages of distorters that later increase in strength. They may also have evolutionary dynamics distinct from strong distorters if they are less prone to having pleiotropic deleterious effects. There is thus a particular need to confront the challenging task of screening for weak and moderate strength distorters in natural populations.

Here we focus on the class of distorters known as meiotic drivers, which manipulate chromosomal segregation to their advantage in individuals that have asymmetric meiosis. In flies and mammals meiotic drive is restricted to females, where only one of the four meiotic products is passed to the oocyte while the remaining three become polar bodies that are not inherited. This creates the unique opportunity for selfish genetic elements to bias their own inclusion into the oocyte resulting in meiotic drive. Centromeres and telomeres are likely hotspots for meiotic drivers to accumulate given their importance in chromosome segregation (Zwick *et al.* 1999). Centromeric sequences are surprisingly poorly conserved; this may be explained if recurrent evolution of meiotic drivers is caused by newly evolved centromeric satellite sequences (Walker 1971). Furthermore if meiotic drivers induce pleiotropic deleterious effects on host fitness, it also explains patterns of adaptive evolution of centromeric proteins as being due to a host response to suppress meiotic drive (Malik 2009).

In Drosophila, telomere-associated sequences also show patterns of rapid evolution. Unlike most organisms which utilize telomerase to extend telomeric sequences, Drosophila telomeres are maintained by three specialized retrotransposons, *HeT-A*, *TART* and *TAHRE* (Mason *et al.* 2008; Pardue *et al.* 2011), collectively known as the HTT elements, that vary widely in sequence and structure among species (Villasante *et al.* 2007; Piñeyro *et al.* 2011). Other sequence classes also point to the evolutionary lability of telomere regions including sub-telomeric repeats (Mefford and Trask 2002; Anderson *et al.* 2008; Mason and Alfredo 2013), sub-telomeric genes (Walter *et al.* 1995; Kern and Begun 2008), and subtelomeric euchromatic sequences (Anderson *et al.* 2008).

As with some centromere proteins, multiple proteins involved in telomere regulation in Drosophila show elevated rates of evolution including the telomere cap proteins Hoap and HipHop, multiple genes in the piRNA pathway, and transposable element (TE) repressors like *Lhr* and *Hmr* (Schmid and Tautz 1997; Gao *et al.* 2010; Lee and Langley 2010; Blumenstiel 2011; Raffa *et al.* 2011; Satyaki *et al.* 2014; Lee *et al.* in press). As mutant alleles of several of these genes produce long telomeres (Khurana *et al.* 2010; Shpiz *et al.* 2012; Satyaki *et al.* 2014), it raises the possibility that telomere length variation causes meiotic drive, with telomere proteins recurrently evolving to counter telomere lengthening

To test this hypothesis, here we sample telomere length variation from a natural population of *D. melanogaster*, focusing on the most abundant telomeric TE, *HeT-A*. We report the discovery of unprecedented variation in telomere length, which provides the material to test whether pairing of extreme telomere length variants in a heterozygous female causes meiotic drive. To do so, we devise a method that sensitively detects deviations from Mendelian segregation using whole-genome sequencing of large pools of individuals. With this method, allele frequencies in BC1 progeny are estimated as a proxy for segregation frequency in F1 heterozygous females. We show that a meiotic drive locus will cause a characteristic decay pattern of allele frequencies in flanking loci. We further present alternative cross schemes to distinguish true meiotic drive effects from viability or developmental effects that also cause allele frequency distortion. We use our method to reject the hypothesis that telomere-length variation causes meiotic drive, but successfully detect a candidate centromere-linked locus that increases its transmission frequency by ~8%, demonstrating that our method offers unprecedented power to detect moderate strength meiotic drivers.

## RESULTS

### Drastic telomere length variation

To evaluate the extent to which telomere length varies in wild populations, we quantified relative *HeT-A* abundance in 182 lines from the *Drosophila* Genetic Reference Panel (DGRP) using qPCR (Mackay *et al.* 2012). For reference we also examined the line *GIII*, which harbors the *Telomere elongation* (*Tel*) mutation and has accumulated long telomeres for over a decade (Siriaco *et al.* 2002). Since *HeT-A* elements are frequently 5’ truncated (George *et al.* 2006), we used primers flanking the 5’ region of the *gag* coding sequence in order to capture mostly full-length copies (Figure 1A). We observed an enormous 288-fold range of *HeT-A* abundance between the highest and lowest lines (DGRP-703 and DGRP-852, respectively). Notably, the top 3 lines (DGRP-161, DGRP-882, and DGRP-703) have higher *HeT-A* abundance than GIII, indicating that extremely long telomeres occur in natural populations, albeit at low frequency. Interestingly, the *HeT-A* quantities distribute along a logarithmic scale (Supplementary Figure 1), arguing that the variation among lines is not simply due to new attachments or deletions, which instead predicts linear changes in abundance.

**Fig 1.**
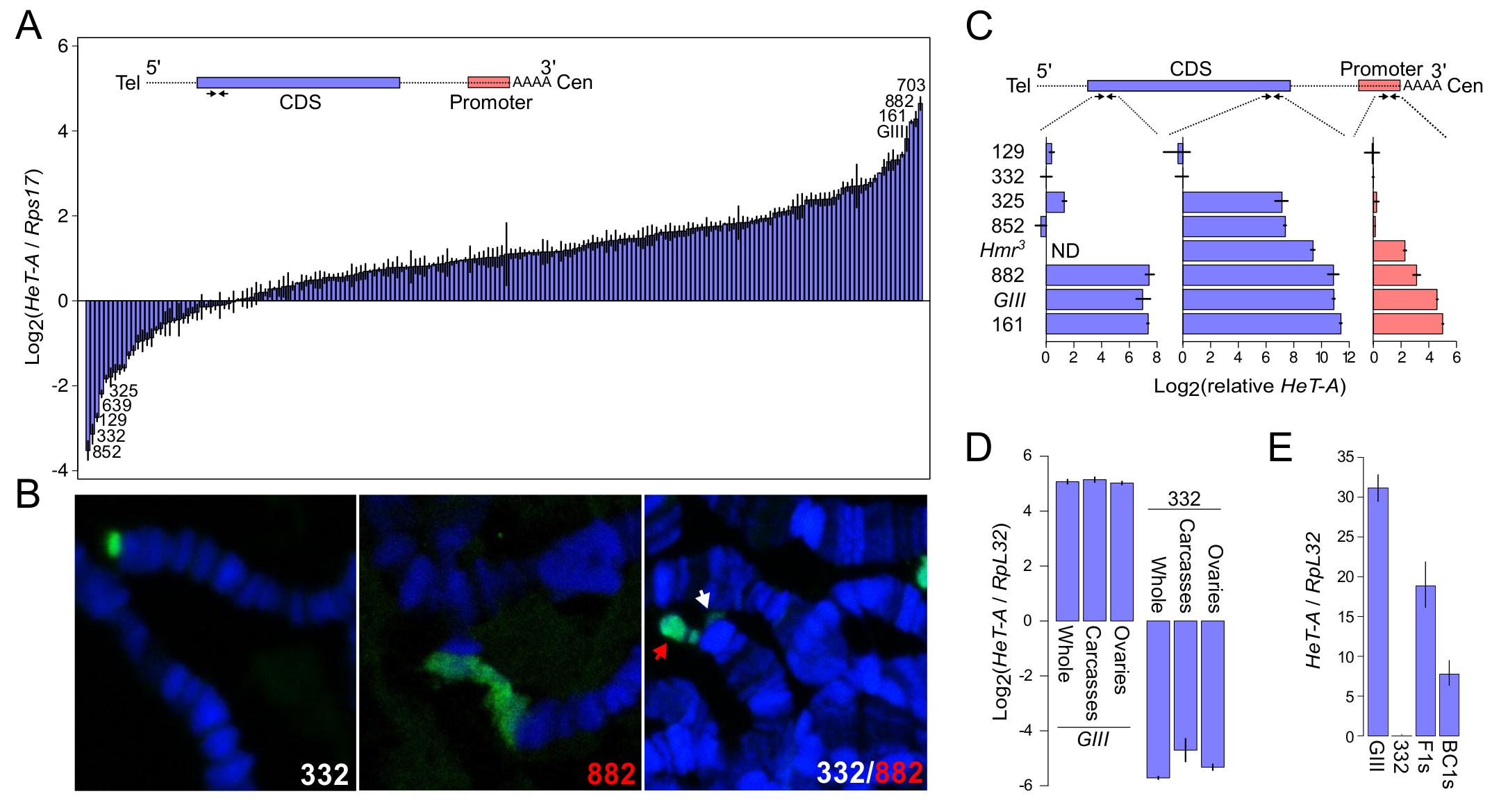
Natural variation in telomere length as assayed by *Het-A* quantities. (A) *Het-A* copy number was measured using qPCR of the DGRP lines as well as the long-telomere strain GIII (Mackay *et al.*, 2012; Siriaco *et al.*, 2002). The primer pair (convergent arrows) targeting the 5’ region is depicted on the schematic of one full length *HeT-A* transcript where the *gag* CDS is colored in blue, the promoter in red, and UTRs are dotted lines. The most extreme lines at each end of the distribution are labeled. Error bars represent the relative error estimated from triplicates. (B) Polytene chromosome spreads of DGRP-332, DGRP-882, and a F1 heterozygote are probed with *Het-A* probe (green). DNA is labeled by DAPI (blue). In the heterozygote, the long allele is labeled by a red arrowhead, and the short allele by an white arrowhead. (C) Relative *HeT-A* quantities were determined at the 3’ CDS and promoter regions as indicated on the schematic in a subset of the DGRP and mutant lines. *HeT-A* quantities at the 5’CDS from Fig. 1A are also re-plotted. Note that different qPCR protocols were used for the two different experiments (see Materials and Methods). Quantities are therefore plotted relative to DGRP-332 so that the three regions can be compared. *Hmr*^*3*^ was not assayed at the 5’CDS (ND). (D) *HeT-A* quantities were measured in whole females, ovaries, and carcasses with ovaries removed in a long (GIII) and a short (DGRP-332) line, using the 3’ CDS primer pair. (E) *HeT-A* quantities were measured in F1s heterozygous for long (GIII) and short (DGRP-332) telomeres and backcross embryos of the F1 females crossed to DGRP-332 males using the 3’ CDS primer pair. Note that the Y-axis here is in linear scale.

We used fluorescence in situ hybridization (FISH) on polytene chromosomes to confirm that the high abundance of *HeT-A* corresponds to long telomeres. We found that the high abundance line DGRP-882 has elongated telomeres marked by *HeT-A* signal, while the low abundance DGRP-332 has markedly less *HeT-A* signal at chromosome ends (Figure 1B). Furthermore, when we crossed the two lines to each other we clearly observed disparate telomere lengths at the tip of a polytene chromosome (Figure 1B). We conclude that *HeT-A* abundance strongly correlates with telomere length, and henceforth refer to high and low abundance lines as having long and short telomeres, respectively.

To further investigate the biological processes underlying the length differences, we quantified abundance at two additional regions of *HeT-A*, the 3’ CDS and the promoter, in a subset of the lines as well as in an additional long-telomere strain, *Hmr*^*3*^ (Satyaki *et al.* 2014). As *HeT-A* is prone to 5’ truncations, differences at the 3’ end likely reflect variation in the insertion rates of both full length and truncated elements. Differences at 5’ regions are expected to be further influenced by rates of incomplete reverse transcription and terminal erosion. We note that because amplification efficiency may differ between primer pairs, it is not possible to directly compare quantities between the three regions. We therefore calculated the respective *HeT-A* abundances relative to the short-telomere line DGRP-332 (Figure 1C). While the relative quantities of *Het-A* are mostly consistent (e.g. the long lines rank the highest in all three regions), there are notable discrepancies. The longest and shortest lines differ by 164-, 2700-, and 33-fold for the promoter, 3’ CDS and 5’ CDS, respectively. These disparities among the three regions suggest that the lines have different proportions of full length versus fragmented elements. For example, the relative *HeT-A* abundances of DGRP-325 and DGRP-852 are similar to DGRP-332 at the 3’ promoter but higher at the 3’ CDS, suggesting that these two lines may have insertion rates similar to DGRP-332 but that they insert longer 5’-truncated elements.

The enormous range observed indicates that telomere length is highly labile. To test the stability of extremely long telomeres, we first compared *HeT-A* abundance in ovaries and carcasses of the extremely long GIII line. We found similar quantities in both, indicating that long telomeres are stable within individuals (Figure 1D). We then tested whether the length is stably transmitted across generations. We crossed GIII to the short DGRP-332 line, reasoning that the discrepant chromosome ends may be particularly unstable in a heterozygous background (Figure 1E). *HeT-A* abundance in these F1s was approximately intermediate to those of the parents. We then backcrossed the F1 females to the DGRP-332 line creating a pool of BC1s and found that *HeT-A* abundance in the BC1s is also intermediate relative to the parents. These results indicate that telomere length is stably inherited across at least two generations.

### Assessing non-Mendelian segregation using pooled sequencing

To test whether drastic telomere length differences cause biased meiotic segregation, we devised a scheme to sample genotype frequencies from whole-genome sequencing of large pools of progeny. We generated heterozygous F1 females by crossing lines with long and short telomeres (P1 and P2, respectively) and then backcrossed the F1s to the P2 parental line (Figure 2A). Under Mendelian segregation, half of the offspring are expected to be heterozygous and the other half homozygous at any given site, assuming that the parents are fully homozygous, thus resulting in an expected ratio of the P1:P2 alleles of 1:3. To minimize the effects of viability and/or developmental differences among individuals of different genotypes, we collected early embryos within a short time window (3-4 hrs after egg laying). Furthermore, to minimize sampling error, we collect large numbers of embryos (between ~1800 and ~8000). After pooling all embryos for each of the crosses, DNA was extracted and sequenced with Illumina to ~30x depth per cross (Table 1, Supplementary Figure 2, see Materials and Methods).

**Fig 2.**
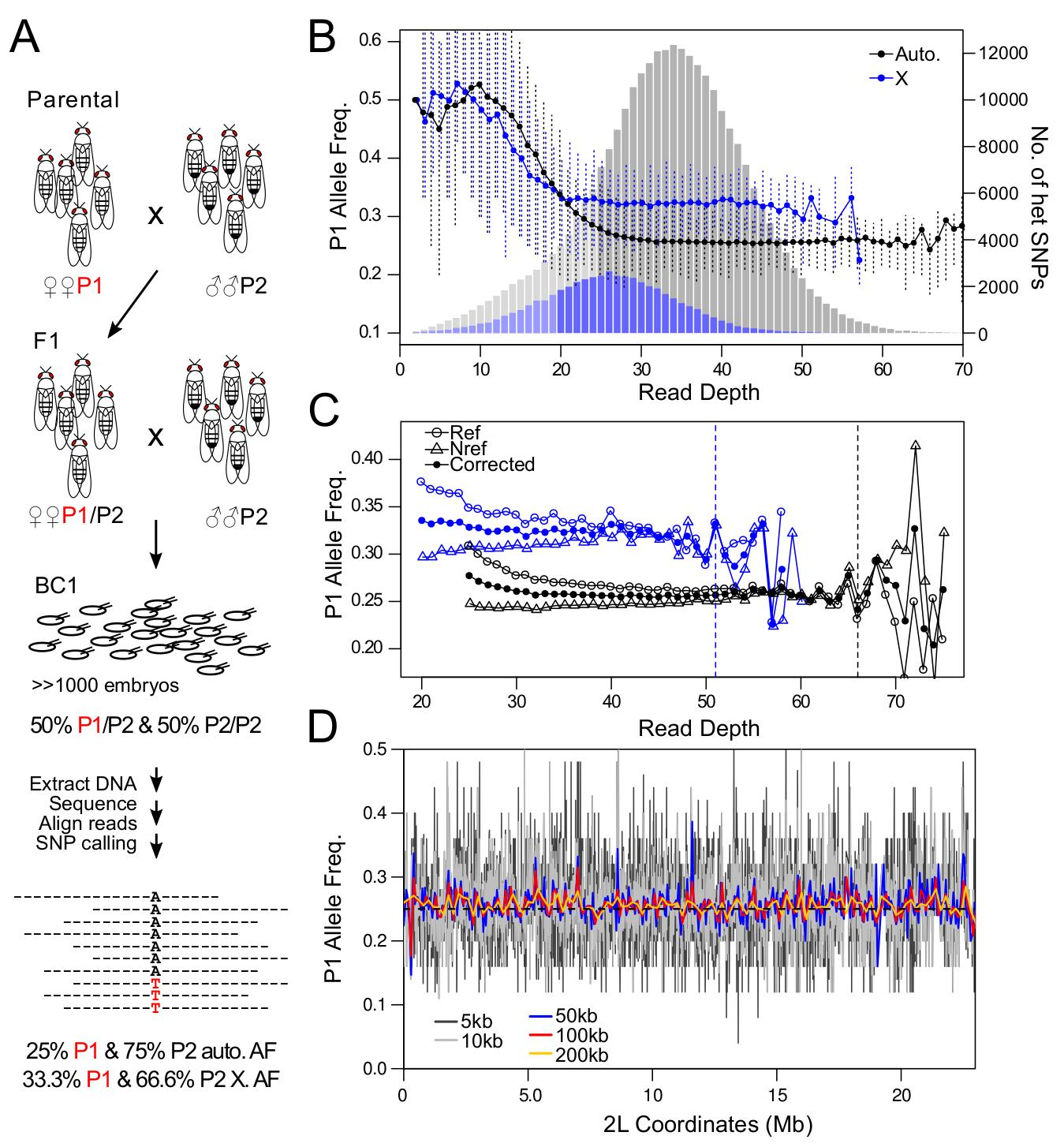
Experimental strategy and statistical considerations for assessing allele frequency by pooled sequencing. (A) Strategy to measure distortion of Mendelian segregation using whole-genome sequencing of pooled embryos. Females (P1) and males (P2) of different telomere lengths are mated to generate F1s. The F1 females are backcrossed to P2 and a large number of 3-4 hr old BC1 embryos collected for sequencing. Heterozygous SNP sites are identified to infer segregation frequency. At the bottom are shown the allele frequencies (AF) expected under Mendelian segregation for autosomal (auto.) and X-linked (X) alleles. (B-D) Analysis of SNPs from the DGRP-882 and DGRP-129 cross. (B) Observed average frequencies of the P1 allele in backcross progeny across heterozygous sites at different read depths are plotted for the autosomal (black line) and X-linked SNPs (blue line). Error-bars delineate the 0.25 and 0.75 quantiles. Underneath is the distribution of autosomal (gray) and X-linked (blue) heterozygous (het) SNPs for each read depth. Bins with lighter shades are sites removed from allele frequency estimation in downstream analyses. (C) The average allele frequency in backcross progeny is plotted when P1 is the reference allele (open circles) and when it is the alternative allele (triangles). The allele frequency after correction for reference is plotted in closed circles. Autosomal and X-linked sites are distinguished by black and blue, respectively. Dotted lines mark the cutoffs for reference allele correction. (D) Aggregating heterozygous sites reduces the sampling noise. Allele frequencies are estimated from simulated reads that have 25% P1 frequency (dotted black lines) and plotted after binning sites at different window sizes (colored lines).

**Table 1.** Summary of pooled sequencing

| P1 | P2 | No. of embryos | No. of reads (million) | No. of heterozygous sites | Autosomes |  | X |  | No. of heterozygous sites after filtering |
| --- | --- | --- | --- | --- | --- | --- | --- | --- | --- |
|  |  |  |  |  | Avg. depth | Avg. P1 freq. | Avg. depth | Avg. P1 freq. |  |
| **GIII** | NC332 | ~3600 | 76.09 | 430,564 | 31.03 | 0.305* | 23.91 | 0.360* | 326,682 |
| **NC882** | NC129 | ~2000 | 89.04 | 418,278 | 33.13 | 0.280* | 25.95 | 0.341* | 343,235 |
| **Hmr<sup>3</sup>** | NC129 | ~1800 | 64.15 | 339,204 | 24.81 | 0.270* | 19.60 | 0.367* | 182,183 |
| NC332 | **NC882** | ~8000 | 88.40 | 440,510 | 33.37 | 0.270* | 25.96 | 0.348* | 362,844 |

\* p value < 1e-6 when compared to expected value of 0.25 for autosomes and 0.33 for X.
Lines with long telomeres are in bold.

To infer allele frequency, we determined heterozygous sites genome-wide in two ways. First, we used GATK to infer SNPs in the pooled sequence reads, from which we then identified heterozygous SNPs. Second, we called SNPs again using GATK in the parental lines from available collections (DGRP lines) or our own sequencing (*Hmr*^*3*^ and GIII), and then identified sites where the parents were homozygous for different nucleotides so that all offspring are expected to be heterozygous. We used only the set of SNPs found in both methods, resulting in ~400,000 heterozygous sites genome-wide for each cross (Table 1). Across these sites, the average frequency of the P1 allele is significantly higher than expected (Table 1). To determine whether this elevation is artifactual, we looked at the P1 frequency across sites of different read depth and found that low-depth sites have markedly higher P1 frequency (Figure 2B). We attribute this to the asymmetric ascertainment bias involved in calling heterozygous sites where the expected allele frequency is 0.25. In a binomial process, a 25% success rate is likely to yield trials where the P1 allele is not sampled, particularly when the number of draws is low (meaning here, at low read depth). This causes a tendency to miss heterozygous sites with low P1 allele frequency, introducing a bias that depends on read depth. To minimize this bias, we removed X-linked and autosomal sites with read depth less than 20 and 25, respectively, reducing the number of heterozygous sites to ~300,000 (Table 1). We note that the *Hmr*^*3*^ x DGRP-129 cross had the largest reduction in heterozygous sites as it had shallower sequencing compared to the other crosses.

Ascertainment bias for the reference allele can also influence inference of allele frequencies. As reference alleles often exist in sizeable haplotype blocks, this bias may result in local deviations from the expected frequencies. We therefore differentiated sites where the P1 allele is the reference from those where P2 allele is the reference. Indeed, the P1 frequency in the former case is significantly elevated (Figure 2C). The difference between the frequencies of reference and non-reference alleles decreases as read depth increases. To correct for this, we estimated the extent of bias for each read depth, and weighted the non-reference alleles accordingly (see Materials and Methods). This correction approach produces similar results to the strategy of aligning to both parental genomes, commonly used for allele-specific gene expression studies (Coolon *et al.* 2012; Satya *et al.* 2012).

While the sampling of alleles might at first be expected to be binomial, there are PCR steps and other features of the experiment that can inflate the sample-to-sample variance (Plagnol *et al.* 2012). In fact, the variance of allele counts is notably greater than binomial (Supplementary Figure 4); instead, we use the beta-binomial process to model the sampling procedure (Cai *et al.* 2012). For these data, variance in allele proportions at both low and high read depths is elevated. This is in part because the left and right tails of the read depth distribution include structural variants in the genome such as deletions and duplications, respectively, which alter both the expected read depth and the allele frequency. We therefore removed sites with greater than twice the average read-depth; along with the lower-bound cutoff mentioned previously, this filter reduces the number of informative sites by less than 25% in the crosses where the average read depth is more than 30x (Table 1, Supplementary Figure 2)

Neighboring sites are expected to have similar patterns of distortion since recombination will rarely break the genetic linkage at small scales. We therefore reasoned that heterozygous sites can be aggregated to increase power and reduce sampling error. To determine an appropriate window size to aggregate heterozygous sites, we generated and processed reads in silico that simulate pooled sequencing of chromosome 2L from a cross between DGRP-882 and DGRP-129 in the absence of any distorting effects. We then estimated allele frequencies along the chromosome by binning filtered and bias-corrected heterozygous sites into nonoverlapping windows of different sizes (Figure 2D), where allele counts of sites within a window are summed. We chose to bin the sites into 200 kb windows, as it substantially reduces the noise in allele frequency estimates along the chromosome. However, a caveat of this aggregation approach is that very closely linked sites are not independently sampled. This issue is most salient for sites within 100 bp, which are almost exclusively sampled together in one sequence read. To determine the extent to which this affects allele frequency and error estimates, we randomly selected one site when two or more are found within 100 bp and removed the rest. We found that this filter has negligible influence on the allele frequency estimates and only slightly increases the error rate (Supplementary Figure 5, as compared to Figure 3).

**Fig 3.**
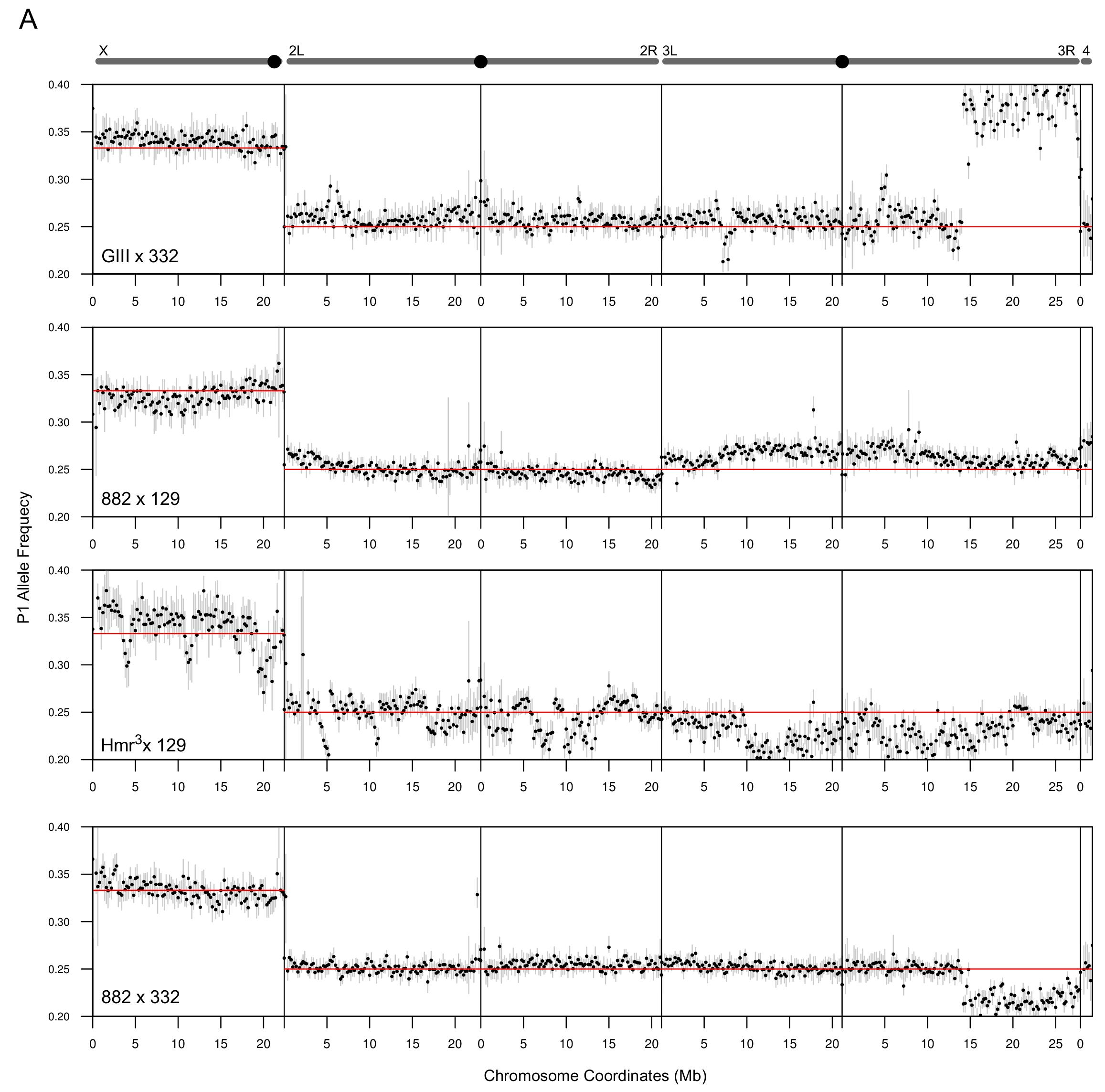
P1 allele frequency estimates across chromosomes. Schematics of chromosomes are shown at the top with centromeres labeled as black circles, except for the 4th chromosome at the far right. For each of the crosses, the frequencies of allele counts at heterozygous sites are averaged in 200kb windows and plotted across all chromosomes. Red horizontal lines mark the Mendelian expectation. Error bars represent 99% confidence intervals which are derived from Monte-Carlo simulations that sum sites with counts randomly generated using the beta-binomial distribution (see Materials and Methods).

### Unaccounted heterozygosity produces sharp fluctuations in allele frequency

The cross between *Hmr*^*3*^ and DGRP-129 yielded the most uneven allele frequencies, with multiple sharp spikes and dips (Figure 3). We attributed these deviations to a combination of low embryo counts and low read depth causing high sampling variance. We also reasoned that a high degree of heterozygosity and structural variants in one or both of the lines might further contribute to the spiky pattern of allele frequencies. To address this we determined the level of heterozygous SNPs within the lines used. The *Hmr*^*3*^ line had a high level of heterozygosity despite having been backcrossed for 6 generations to an inbred strain (Figure 4); DGRP-129, on the other hand is relatively free of heterozygosity.

**Fig 4.**
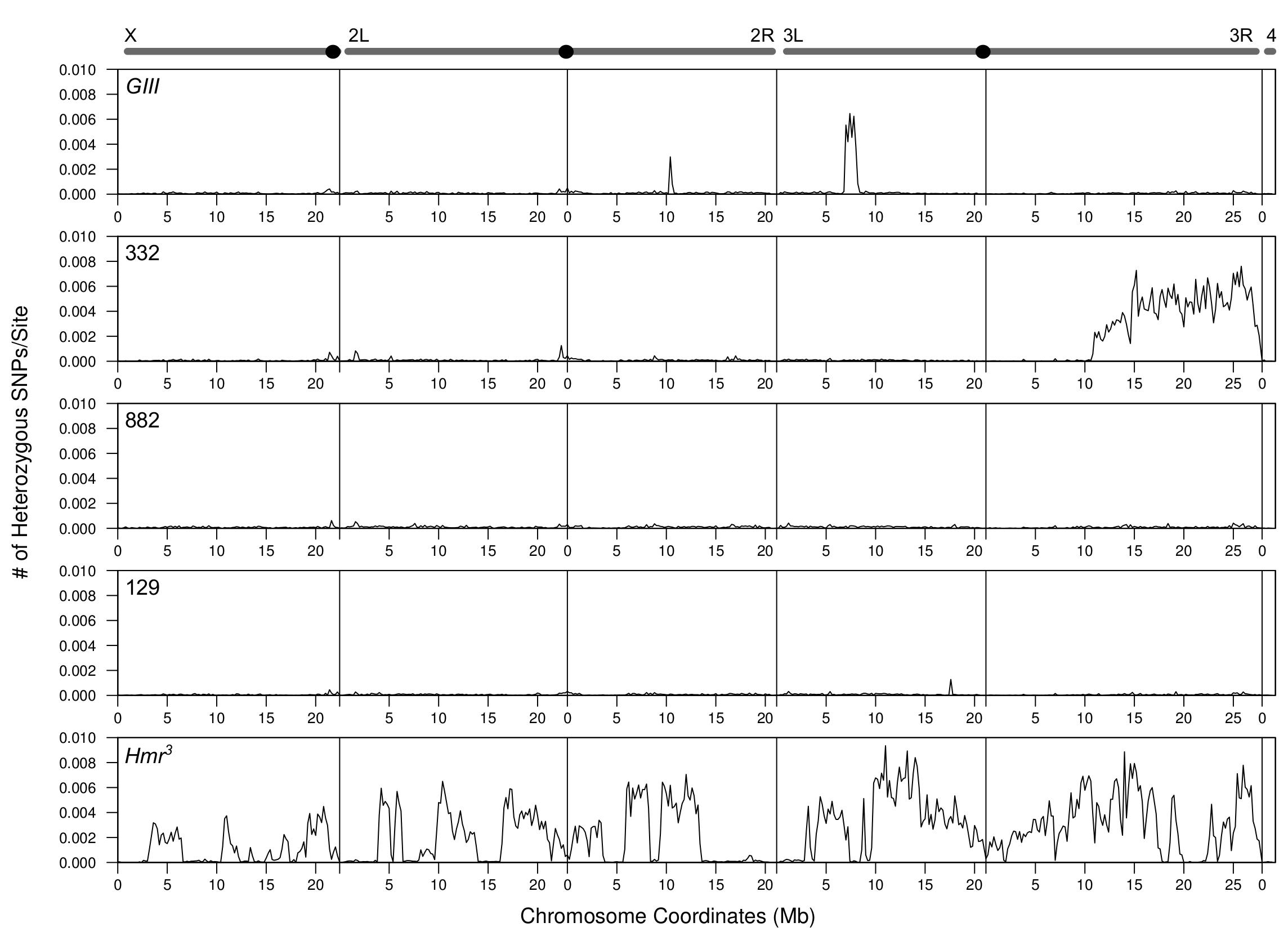
Frequency of heterozygous sites of parental lines. For each of the parental line, the frequency of heterozygous sites per base is plotted across the chromosomes in 200kb windows.

On chromosome 3R of the GIII x DGRP-332 and DGRP-882 x DGRP-332 crosses, we detected distortion spanning over half of the chromosome, starting at around 15 Mb; the allele frequency increased to an average of 0.384 and dropped to 0.217, respectively. As DGRP-332 was used in both crosses and the direction of distortion was opposite in the two crosses, we reasoned that the non-Mendelian ratios are due to a large structural variation or polymorphism in the DGRP-332 line. We indeed found that the distal half of chromosome 3R has a very high level of polymorphism (Figure 4). Using PCR across indel polymorphisms, we further confirmed that DGRP-332 is heterozygous at three sites in the distal half of chromosome 3R (see Materials and Methods). The remaining lines have very low levels of heterozygosity across the chromosomes, with the exception of two small regions in GIII. These results demonstrate that despite our filtering strategy to only select homozygous sites in the parental lines, heterozygous sites remain in some lines and can dramatically distort allele frequencies, especially if they are in large blocks.

### Meiotic drive produces distortions with a characteristic decay pattern

While structural variants and heterozygosity create deviations in allele frequency with sharp boundaries, meiotic drive produces distortion signals that should be qualitatively distinct. As our experiments are mapping across one generation of recombination, we expect that a driver will produce a broad peak of distortion as it will also elevate linked loci. For a driver that is not contained within an inversion, as recombination breaks the linkage between it and flanking sites, distortion is expected to attenuate with genetic distance, vanishing by 50 cM as the driver and SNPs become fully unlinked. Drivers within inversions are expected to show a broader peak, with the shape of the signal further dependent on whether or not recombination is fully suppressed within the inversion.

We noticed slight elevations of the P1 frequency in the DGRP-882 x DGRP-129 cross at two regions, both of which show a gradual decay. The first is at the telomere of 2L, where the elevated P1 allele frequency attenuates toward the centromere (Figure 5A), and the second is at the centromere of chromosome 3, where the elevated allele frequency decreases on both arms distally (Figure 5B). To determine whether the observed decay of the signals is consistent with recombination breaking the linkage between the drive locus and distal loci, we simulated the decay of the signals based on published estimates of recombination rates across the chromosome arms (Fiston-Lavier *et al.* 2010), and also determined the magnitude of deviation that best fits the recombination rates (Supplementary Figure 6C and D, see Materials and Methods). We estimate that the distortion on 2L increases the allele frequency to 0.260 (± 0.00448, 95% confidence interval), equating to a 4.2% increase in transmission frequency. The observed allele frequency decays faster than the simulated decay, as the former drops to the Mendelian expectation at around 10 Mb while the latter fully attenuates at ~16 Mb. One possible explanation is that this discrepancy reflects variation in recombination rate among lines. The centromere distortion on chromosome 3 elevates the allele frequency to 0.271 (± 0.00415) equating to a 8.4% increase, and the observed decay fits the simulated decay strikingly well. Because DGRP-882 shows no effect when crossed to DGRP-332, we suspect that the observed deviation is caused by polymorphisms present only between the DGRP-882 and DGRP-129 stocks. Since only one chromosome end out of all the crosses between long and short telomere lines displayed deviation consistent with meiotic drive, we conclude that extreme telomere length differences are insufficient to cause biased segregation.

**Fig 5.**
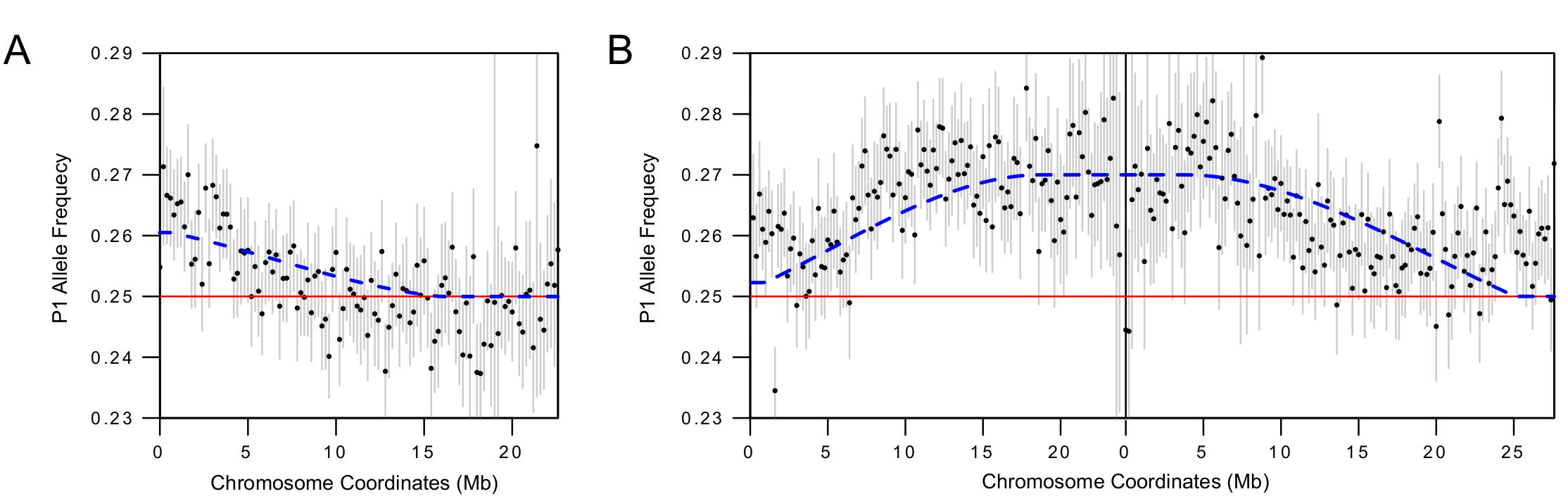
Allele frequency decay due to recombination. (A-B) Magnified views of chromosomes 2L (A) and 3 (B) from the DGRP-882 X DGRP-129 cross. The blue dotted line indicates the expected decline in signal of distortion for a telomeric (A) or centromeric (B) drive locus, based on genome-wide recombination rate estimates.

### Crossing schemes to distinguish meiotic drive from developmental effects identifies a candidate centromere-linked driver

While meiotic drive causes distortions to Mendelian ratios and by extension allele frequencies, developmental effects arising from polymorphism between the parental lines can also cause allele-frequency deviations. The progeny pool contains P1/P2 heterozygotes and P2/P2 homozygotes. These genotypes may have phenotypic differences in viability, embryo size, cell number, and/or growth rate, which we collectively term developmental effects, that can produce unequal representation of the two genotypes resulting in P1 allele frequencies that deviate from 0.25. To distinguish whether the distortion signals we found in the DGRP-882 x DGRP-129 cross are consistent with meiotic drive versus developmental effects, we altered the crossing scheme in two ways. First, we performed an alternative backcross of the F1s to P1 males instead of P2 males, producing a pool of P1/P1 homozygotes and P1/P2 heterozygotes (Figure 6A, left). If meiotic drive is causing the deviation observed in the initial backcross, it should also be observed in this alternative backcross because P1 transmission will be increased in the heterozygous mother irrespective of the sire genotype (Figure 6B). Second, we performed a reciprocal cross using F1 heterozygous males back-crossed to P1 females (Figure 6A, right). This cross should not show any deviation because meiotic drive only occurs during meiosis of heterozygous females. The combination of the three crosses also allows us to distinguish among a range of developmental effects. For example, if heterozygotes are in general more fit than homozygotes due to heterosis, this would cause elevations of P1 frequencies in the initial backcross as observed in Figures 3B and 3C, because P1/P2 heterozygotes are being pooled along with P2/P2 homozygotes. But in the alternative backcross and reciprocal cross, the P1 frequencies of the pools will be lower than expected, not higher, since P1/P2 heterozygotes will be more fit than P1/P1 homozygotes (Figure 6B). Finally, additive viability effects will equally inflate P1 transmission frequencies in all 3 crosses (Figure 6B). Because recombination does not occur in male Drosophila, both meiotic drive and developmental effects will cause deviations in the reciprocal cross across the entire chromosome on which they occur. We also note that adults were collected for these experiments instead of embryos. This was done largely because of the effort of collecting large numbers of staged embryos, but we also reasoned that significant deviations caused by meiotic drive are likely to persist irrespective of developmental time and life stage.

**Fig 6.**
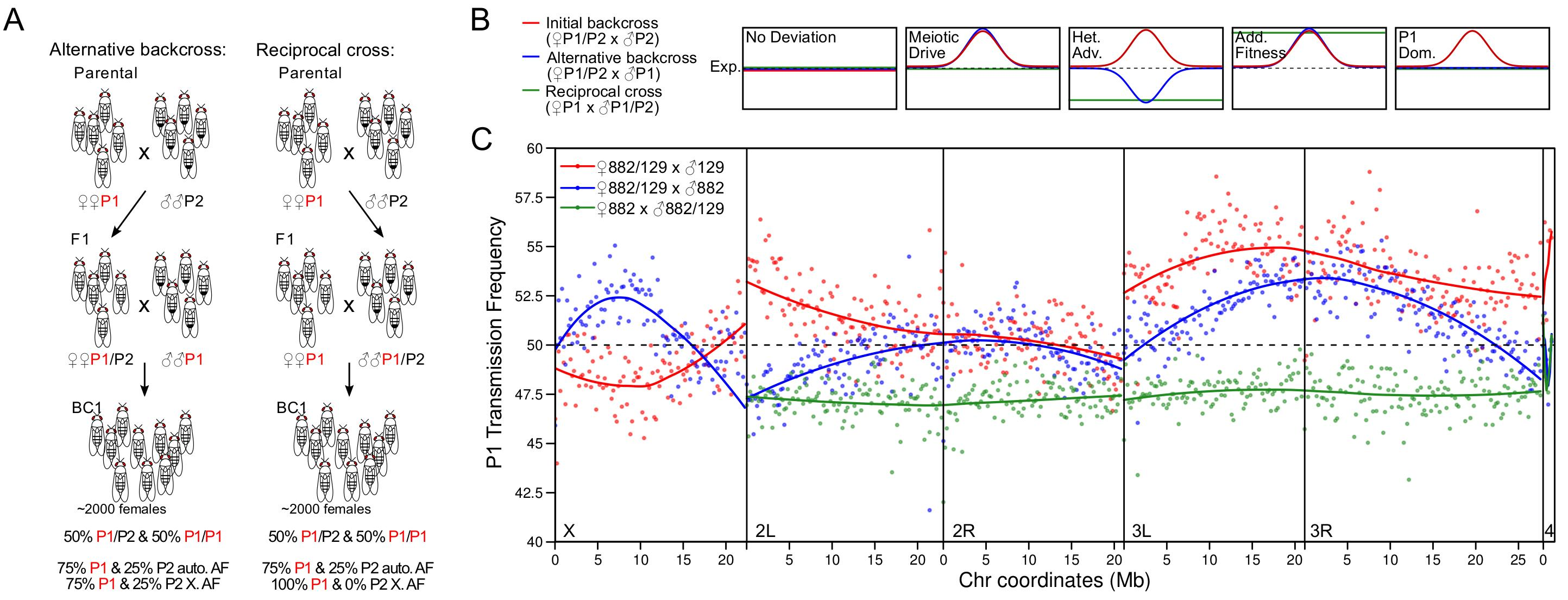
Crosses to distinguish developmental effects from meiotic drive. (A) Two cross schemes to determine whether the deviations observed with the initial cross scheme (Fig 2A) are true meiotic drive effects as opposed to developmental effects. At the bottom are shown the genotype frequencies above the autosomal allele frequencies (AF) expected under Mendelian segregation. Note that only adult females were collected. (B) Patterns of deviation of P1 transmission frequency from the Mendelian expectation (dotted lines) depending on the cross scheme (colored lines) are depicted for meiotic drive, heterozygous advantage (Het. Adv), and P1-allele-specific fitness advantage that is additive (Add. Fit.) or dominant (P1 Dom). (C) The transmission frequency of the P1 allele (DGRP-882), averaged across 200 kb windows, is plotted for the initial cross from Fig. 3A (red), the alternative backcross (blue), and the reciprocal cross (green). The curves for each cross scheme and chromosome were fitted using local regression (LOESS).

To facilitate comparison between the different crossing schemes, we transformed the allele frequency estimates into transmission rate of the P1 allele by removing the contribution from the parental line used in the backcross, such that non-Mendelian transmission will cause deviations from 50% (see Materials and Methods). In the alternative backcross, the elevation in P1 previously linked to the telomere of 2L is instead a depression, indicating that more robust heterozygotes instead of meiotic drive is likely the cause of the deviation (Figure 6C). The reciprocal cross also supports this interpretation because it gives a uniform deviation of ~47.5%, which is very similar in magnitude to that observed at the 2L telomere in the alternative backcross. Similarly, the slight depression on the X in the initial cross became an elevation in the alternative back-cross.

Strikingly however, we see a persistence of a similar elevation in the alternative backcross for the elevation linked to the centromere of chromosome 3. Moreover, this elevation is not observed in the reciprocal cross, indicating that distortion only occurs during female meiosis and is thus consistent with meiotic drive. There are clearly some differences between the two backcrosses. The distal end of chromosome 3L shows distortion in the original backcross to P2 but not in the backcross to P1, which is consistent with a P1 dominance effect. This effect as well as the different degrees of signal attenuation at the distal end of chromosome 3R may explain the apparent shift in the centromere region peak between the two backcrosses. In the reciprocal cross, the transmission frequency on chromosome 3 is consistently lower than the Mendelian expectation. While the absence of signal attenuation makes it impossible to locate the source of the deviation, the P1 transmission frequency of ~47.5% is closely similar to that seen at the distal end of 3R in the alternative backcross, suggesting that both reflect heterozygote advantage.

## DISCUSSION

### Identification of biased segregation using whole-genome sequencing

Most known cases of non-Mendelian chromosome segregation involve loci of large effect (Fishman and Willis 2005; Larracuente and Presgraves 2012). The difficulty in genotyping large numbers of individuals to minimize sampling error has limited the ability to identify weak drivers, which are expected to be more frequently found in natural populations at intermediate frequencies because they are slower to reach fixation and less likely to have strongly deleterious pleiotropic effects (Thomson and Feldman 1976).

Our strategy of sequencing pools of individuals has several advantages that allow us to identify candidate loci of weak drivers. First, by assaying allele frequencies from pools, we are able to sample thousands of individuals, thereby minimizing sampling error and increasing sensitivity to detect distortion. Second, by using whole-genome sequencing, we can search for drive loci genome-wide and map them (at low resolution), as opposed to targeted genotyping which requires prior knowledge of the location of the driver. Third, the large number of informative heterozygous sites genome-wide provides information for error and bias correction. Lastly, the contiguous windows of allele frequencies across chromosomes allow us to detect and fully visualize the attenuation of distortion as predicted using estimates of recombination rate.

Similar strategies have been applied on sperm pools of hybrid male mice (Corbett-Detig *et al.* 2015; Bélanger *et al.* 2016). In the first study, direct sampling of gametes as opposed to F1s has the major advantage of eliminating viability effects. This strategy may be applicable on sperm isolates in flies (Dorus *et al.* 2006), but would not work for *Drosophila* oocytes as they maintain their polar bodies until after fertilization. Instead, generating haploid embryos by eliminating the paternal genome might allow for direct sampling of female meiotic segregation (Langley *et al.* 2011). However, the efficacy will depend on how uniformly the haploid embryos arrest and the number of gynogenetic haploid escapers. In the second study, genotype-by-sequencing, a form of reduced representation sequencing, was used to identify loci with distorted segregation rates (potentially caused by a range of mechanisms). By reducing the number of loci sampled, this approach offers deeper sequencing per site and therefore can accommodate more samples per sequencing run.

Given that our approach tracks chromosome-wide allele frequency, we can detect minor distortions that reflect increases as little as 5% in segregation frequency (Supplementary Figure 7). We stress that the expectation of gradual signal attenuation due to recombination plays a pivotal role in qualitative assessment of candidate distortion loci. Significant deviation from the expected Mendelian ratio can be caused by structural variants like indels and duplications or by unaccounted residual heterozygosity in the parental lines (Figures 3, 4). Both will produce alterations to the expected allele frequency, but the resulting distortion will be local, and, importantly, will have sharp edges rather than attenuating gradually.

The observation of decay in allele-frequency distortion cannot however distinguish whether the deviation is caused by meiotic drive or developmental differences among genotypes. The latter can result from differences in growth rate, body size, or viability, all of which will produce an unequal contribution of DNA and therefore distortion at the causal loci as well as a signal of decay due to recombination. We therefore used additional crosses to differentiate between meiotic drive and different types of developmental effects. By backcrossing to the P1 parent instead of P2 and by performing a reciprocal cross with heterozygous males, we successfully showed that several deviations observed in the initial cross with DGRP strains DGRP-129 and DGRP-882 are likely caused by heterozygote advantage. The additional crosses can also distinguish between additive, dominant, and recessive effects. For example, an additive effect will manifest as deviations in the same direction for both backcrosses, but P1 dominance will only cause deviation in the backcross to P2 males (Fig. 6B).

While the use of the reciprocal cross is particularly diagnostic of female-specific meiotic drive, there are several caveats. First, it cannot detect X-linked deviations as it is impossible to have a pair of Xs in males. Second, the absence of male recombination in Drosophila means that any fitness effect will cause deviations of the entire chromosome as observed in Fig. 6C. Third, maternal and paternal effects can potentially cause developmental effects that are specific to either crosses with female heterozygotes or to the reciprocal cross. In particular, maternal effects that result in an additive fitness increase for P1 alleles would mimic the signal of meiotic drive. Distinguishing these possibilities will require comprehensively genotyping individual embryos. Nevertheless, we establish here that sequencing of pools of progeny is an effective and sensitive method to identify candidate meiotic drivers of moderate strength and to reveal their approximate location in order to design further mapping studies.

### Massive *HeT-A* and telomere length variation

We assessed natural variation in telomere length across the DGRP lines by quantifying relative *HeT-A* copy number using qPCR. These data quantitate total *HeT-A* abundance per line and thus do not provide any information on possible variation among the different telomeres within lines. As most *HeT-A* elements are truncated at the 5’ end, quantification of the 5’ ends likely represents full-length elements. Surprisingly, we observed an enormous 288-fold range between the longest and shortest lines. The abundance is distributed on a logarithmic scale indicating that telomere length increases non-linearly. This is inconsistent with a simple model where *HeT-A* additions occur at a constant rate, and suggests instead that high abundance arrays are more likely to have a higher rate of gain. The observed distribution can result from attachment events, if the rate of addition increases as more elements accumulate, increasing the copy number of active elements. Alternatively, unequal crossing-over can increase and decrease large blocks of *HeT-A* arrays creating a large range of sizes. In addition, the probability of crossing-over is expected to increase as telomeres become longer.

Determining a precise quantitative estimate of telomere length in Drosophila is more challenging than in other eukaryotes because its telomeric repeat sequences are much more complex. Assuming that the lowest line has the minimum of 1 full-length *HeT-A*, the mean across the DGRP lines is 34 copies, and the *y w* line used as reference has 25 copies. These numbers are similar to that of the genome reference line (*y*^*1*^*; cn*^*1*^ *bw*^*1*^ *sp*^*1*^) that was independently estimated to have approximately 29 and 7 full-length copies of *HeT-A* and *TART*, respectively (George *et al.* 2006), which amounts to 365 kb of *HeT-A* and *TART* sequences in total and 45.6 kb per chromosome end. Based on these numbers, the longest lines we assayed would have over 3.6 Mb of telomeric sequence equating to an average of ~450 kb per chromosome end. The shortest line would have only 12 kb in total and 1.5 kb per chromosome end, but given the large number of truncations, the actual sizes of the telomeres in short lines are likely larger. Because we only sampled *HeT-A*, we cannot rule out the possibility that *TART* and *Tahre* might compensate for low *HeT-A* numbers. However, this is unlikely, as surveys of *HeT-A* and *TART* quantities across several stocks, including GIII, found that increases in *Het-A* abundance are accompanied by increases in *TART* abundance (Siriaco *et al.* 2002).

The range of estimated sizes that we detected contrasts with organisms that utilize the canonical telomerase mechanism of telomere protection. Telomere lengths have been estimated to be between 5-10 kb in humans, ~1 kb to 9.3 kb in *Arabidopsis thaliana*, ~150 to 3500 bp in *C. elegans*, and only <100 to 500 bp in yeast (Vasa-Nicotera *et al.* 2005; Liti *et al.* 2009; Thompson *et al.* 2013; Fulcher *et al.* 2015; Cook *et al.* 2016). Furthermore, even interspecific differences such as that between mouse species pale in comparison (Zhu *et al.* 1998). These differences may be due to the retrotransposon-based mechanism of telomere evolution found in Drosophila. We suspect that the evolution of this mechanism allows telomere length to be more labile in flies. Interestingly, this lability may also present challenges for proper regulation, thereby driving the rapid evolution of multiple components of the capping complex. More stable telomeres, in contrast, may lead to regulation that is highly conserved, as exemplified by the CST and shelterin complexes in yeast and humans, respectively (Raffa *et al.* 2011).

### Potential genomic elements causing meiotic drive

By generating F1s heterozygous for long and short telomeres, we tested the hypothesis that telomere length biases chromosome segregation in meiosis. However, among 4 crosses tested, no chromosome end in any cross showed allele frequency distortion consistent with meiotic drive. Thus, we conclude that large telomere-length differences, at least as assayed by *HeT-A* quantities, are insufficient to cause meiotic drive.

To our surprise, however, we identified a candidate driver in the cross between DGRP-882 and DGRP-129 in the region spanning the centromere of chromosome 3. This candidate driver increases the transmission of the DGRP-882 allele by ~8%, which corresponds to a 54:46 segregation ratio. Notably, we did not detect distortion in the crosses between DGRP-882 and DGRP-332 or between *Hmr*^*3*^ and DGRP-129. These results suggest that the cause of distortion reflects polymorphism(s) between DGRP-882 and DGRP-129 that are not variable between the other strain pairs.

The potential for centromeres to accumulate meiotic drivers has been long recognized (Walker 1971; Pardo-Manuel de Villena and Sapienza 2001; Malik 2009). One clear example comes from the centromere-linked locus *D* in *Mimulus* that causes near mono-allelic segregation in interspecific hybrids and 58:42 segregation ratio in intraspecific crosses (Fishman and Saunders 2008). Another is Robertsonian fusions of mouse chromosomes that create metacentric centromeres that segregate into the pronucleus more frequently than the telocentric counterparts (Chmátal *et al.* 2014). This effect is mediated by increased attachment of kinetochore proteins at the “stronger” centromere. Differential amounts of repetitive DNA in flies may similarly create centromeres of different strengths. One of the most abundant satellites in *D. melanogaster,* AACATAAGAT, is a candidate for the effect we observed, as it is located in the pericentromeric regions of 2L and 3L (Lohe *et al.* 1993). We also recently characterized two new satellites mapping near the centromeres of chromosomes 2 and 3; interestingly, they are population-specific with a global distribution that is inconsistent with the neutral expectation (Wei *et al.* 2014). The potential effects of these satellites on meiotic segregation can be tested using the method presented here.

## MATERIALS AND METHODS

### Estimating *HeT-A* quantities with qPCR

For quantification of the DGRP lines at the 5’ CDS in Fig. 1A, approximately ten adult flies of mixed sex from each DGRP line were collected from stock vials and flash frozen in liquid nitrogen. Flies were homogenized by using beads and purification steps as per the manufacturer’s protocol using Agencourt DNAdvance Genomic DNA Isolation Kit (Beckman Coulter). DNA isolation steps were handled by Biomek 4000 Liquid Handling System (Beckman Coulter robotic system). After purification using columns, DNA was eluted in 50 ul sterile water, concentration was estimated by using a NanoDrop 2000 (Thermo Scientific) and diluted to a concentration of 10 ng/µl using sterile water. 5′TTGTCTTCTCCTCCGTCCACC3′ (forward) and 5′GAGCTGAGATTTTTCTCTATGCTACTG3′ (reverse) were used for qPCR. The quantifications were normalized to quantities of RpS17 amplified by the primers 5′AAGCGCATCTGCGAGGAG3′ (forward) and 5′CCTCCTCCTGCAACTTGATG3′ (reverse). Real-time PCR was run using ABI Prism 7900 HT Sequence detection system (Applied Biosystems). Each DNA sample was run in triplicate to estimate average Ct values. Mean and standard deviation values of the three replicate reactions were used to estimate the telomere length of each line.

For quantification of the 3’ CDS and promoter regions in Figure 1C, we collected ~10 1-3 day-old females. DNA was extracted from carcasses with ovaries removed using Puregene Core Kit, and concentration was measured by NanoDrop. Primer sequences for the two regions of *HeT-A* and for RpL32 (rp49) used for normalization were from (Klenov *et al.* 2007).

### Fluorescent *in situ* hybridization on polytene chromosomes

Third instar larvae were dissected in 0.7% NaCl. The salivary glands were separated from the brain and imaginal disks and fixed in freshly made 1.84% paraformaldehyde/45% glacial acetic acid for five min on siliconized coverslips. A frosted glass slide was then applied onto the coverslip and gentle pressure applied to dissociate the cells. The slides were then submerged in liquid nitrogen for at least 10 min with the coverslips then quickly removed with a blade and the slides washed and dehydrated with 95% EtOH. Prior to hybridization the slides were treated with 2X SSC at 70°C for 30 min, followed by dehydration with 95% EtOH at room temperature for 10 min and air-drying for 5 min. The slides were then submerged in 0.07N NaOH for 3 min, followed by dehydration and air-drying again. Remaining steps of hybridization and washing were as in reference (Larracuente and Ferree 2015). For the *HeT-A* probe, a 105 bp HeT-A fragment was amplified using the primer pair CGCAAAGACATCTGGAGGACTACC/ TGCCGACCTGCTTGGTATTG and cloned. The vector was used as a template to generate probes using the PCR Dig Probe Synthesis Kit (Roche). Imaging was carried out with a Zeiss Confocal Microscope and images processed using Zeiss Zen software.

### *Drosophila* stocks and crosses, embryo collection, and sequencing

The long-telomere stocks GIII and *Hmr*^*3*^ are described in (Siriaco *et al.* 2002) and (Satyaki *et al.* 2014), respectively. Identity of DGRP lines used for telomere drive crosses was tested using a subset of RFLPs described in (Mackay *et al.* 2012). All crosses were performed at 25°C. F1 females were generated by setting 3 vials each with ~25 virgin females and ~25 males, and transferring flies to fresh vials every 1-2 days for approximately 1 week. We aimed to have approximately 400 males and 400 virgin females aged 3-7 days as parents for each biological replicate to generate BC1 embryos; in order to have sufficient parents female age ranged from 0-11 days. All vials containing F1 females were kept for several days and monitored for larvae to ensure that they contained only virgins.

Crosses were set with approximately 200 parents in a vial for one or two days, then transferred to an egg-collection cup containing a grape-juice/agar plate supplemented with yeast paste, and kept overnight in the dark. The next day, fresh plates were changed every hour and then aged for 3 hours after collecting. The approximate number of embryos was recorded, embryos rinsed in dH_2_0 and then dechorionated in 50% bleach for 2 minutes. Embryos were examined under a microscope and bleaching continued as necessary. Any larvae and late stage embryos were removed and embryos rinsed with dH_2_0. Embryos were then suspended in PBT and transferred to 1.5 ml eppendorf tubes, excess PBT removed, and vials frozen in liquid nitrogen and stored at −80°C. We used the DNAeasy Kit to extract DNA from embryo pools. Libraries were generated using the TruSeq Kit (Illumina) and sequenced on 2 lanes of HiSeq 2500 Rapid Run (100 bp single-end reads).

For the adult pools, 5-day-old sexed flies were collected and stored in −80°C. ~2000 frozen flies were pulverized in liquid nitrogen with a mortar and pestle and then lysed overnight at 65°C in 30 ml of cell lysis buffer from the Gentra Puregene Tissue Kit for DNA Extraction (Cat No. 158667), followed by incubation with 150 ul of Proteinase K at 55°C for 3 hrs. 300 ul of the lysate was then processed following the manufacturer’s instructions. Illumina libraries were then generated using the TruSeq Kit and sequenced on 1 lane using NextSeq (75 bp single-end reads). All sequences are available from the NCBI SRA under BioProject accession PRJNA350856.

### Processing sequences and identification of heterozygous sites

Sequences were aligned to *D. melanogaster* reference (r5.46) using BWA on standard settings. For genome coverage, we used genomecov in BEDTools (Quinlan and Hall 2010). Using Samtools (Li *et al.* 2009), we sorted the reads, removed PCR duplicates, and merged all samples including both the parental and cross sequences into one file. We applied the GATK package following the recommended practices (McKenna *et al.* 2010; DePristo *et al.* 2011) up to HaplotypeCaller (see https://www.broadinstitute.org/gatk/): we used sequence RealignerTargetCreator (without input of known targets) and IndelRealigner on default settings and HaplotypeCaller with -stand_call_conf of 30. The resulting vcf file, which contains all samples, was then separated into individuals files for each parental and cross sample. For the parental files, only homozygous non-reference SNPs with genotype quality >20 (PHRED scale) were kept. Then, to determine expected heterozygous sites from the parental lines of each cross, we identified sites where only one of the two parents is homozygous for the non-reference allele. To determine heterozygous sites from the crosses, we filtered for single nucleotide heterozygous sites with genotype quality >20 and are biallelic. This set of observed sites was then polarized using the expected parental set, such that the parent of origin of the reference and non-reference alleles is determined. Only heterozygous sites found in both sets were used for allele frequency quantification.

### Allele frequency quantification and reference bias correction

Read counts for each of the two alleles at heterozygous sites were inferred from the AD field in the vcf files, which contains the counts for the reference and non-reference alleles which were polarized.

For each read depth bin *i*, we determined the average P1 allele frequency when it is the reference (*FreqR*_*i*_) and when it is the non-reference allele (*FreqN*_*i*_). The observed allele frequencies can be modeled as:

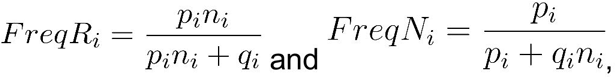

where *n* is the factor to which the reference allele is elevated from the non-reference allele. *p* and *q* are the P1 and P2 counts in the absence of bias, respectively. Based on the two equations, it can be determined that:

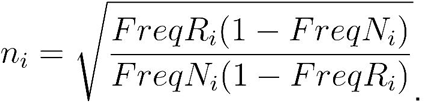

The observed P1 and P2 counts are divided by *n* to determine *p* and *q*, when they are the respective reference alleles. This correction was applied to sites with read depth less than twice the average read depth.

Across multiple sites within a 200 kb window, the counts of alleles from the same parent are summed.

### Over-dispersion and error modeling with beta-binomial sampling

For each read depth bin, *n*, a binomial distribution was fitted with a mean of *np*_*n*_, where *p* is the average P1 frequency at the read depth. To fit the beta-binomial distribution, we used the R package vgam to infer the mean, *np*_*n*_, and a shape parameter, *ρ* estimated from all P1 frequencies (Yee 2015). Beta-binomial distributions were then generated for each read depth with mean of *np*_*n*_ and *ρ*.

To determine the confidence interval of each window based on the beta-binomial sampling process, for a given window with *m* informative heterozygous sites, the aggregated counts of the window are 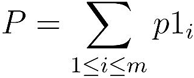 and 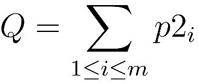 for the P1 and P2 alleles, respectively, where *p*1 and *p*2 are the respective counts at individual sites.

For each window, we iterate the function 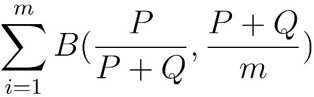 10000 times, where *B*(*a*, *n*) generates a random beta-binomial draw with a probability of *a* and sample size of *n* with the *ρ* shape parameter as determined by VGAM. This generates a distribution for *P*, from which the 99% confidence interval was determined.

### Simulation of pooled sequencing reads with drivers of different strengths

We first used the GATK tool FastaAlternateReferenceMaker to generate pseudo genomes (fasta files) of DGRP-882 and DGRP-129 using their respective vcf files. We then simulated reads from chromosome 2L at different read-depths to simulate different strength of segregation distortion with a combined read depth of 28x using the short read simulator ART (Huang *et al.* 2012). E.g. For Mendelian segregation with a 25:75 ratio of P1:P2, we generated reads at 7x and 21x from DGRP-882 and DGRP-129, respectively. For 10% drive at 27.5:72.5 ratio, we generated reads at 7.7x and 20.3x from the two. The reads were then processed as described above.

### Recombination rate estimates and distortion signal decay

The formulae for map distance measured in cM as a function of physical distance measured in Mb for each chromosome were taken from the *Drosophila melanogaster* recombination rate calculator (Fiston-Lavier *et al.* 2010). The genetic distance (*G*) from the distortion loci was calculated in 200 kb windows, with windows >50 cM set to 50 cM. For each window (*k*) away from the distortion loci on the P1 chromosome, the proportion (*p*) of oocytes carrying the P1 allele can be calculated as: 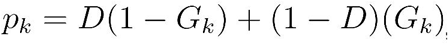, where ? is the proportion of P1 alleles at the distortion locus in the oocyte pool, calculated as twice the P1 allele frequency at the distortion loci on autosomes and 3/2 for loci on the X. The P1 allele frequency at the driver was determined using a least-squares approach, where multiple decay curves were generated with incremental changes (0.0001) in P1 allele frequencies; the P1 allele frequency/curve of best fit was selected. The left and right sides of the addition correspond to the proportions of gametes where the P1 alleles are linked with the distortion locus (i.e. no recombination happened in between) and the P1 alleles are not linked with the distortion locus (i.e. recombination happened in between), respectively. The expected P1 allele frequency is then 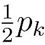 for autosomal sites and 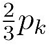 for X-linked sites. Confidence intervals were calculated as per (Geer 2014).

### Transmission rate calculation

To obtain P1 transmission rate from allele frequency for the initial back-cross where we sampled mixed-sex embryos we divided the X and autosomal allele frequencies by 0.667 and 0.5, respectively. For the alternative back-cross and reciprocal cross, we subtracted 0.5 from the allele frequency and divided the difference by 0.5 (these crosses only sampled females).

### PCR genotyping

We identified 20-45 bp indel polymorphisms from the VCF files that distinguish DGRP-332 from GIII across chromosome 3R. We designed two PCR primer pairs flanking indels outside the distortion region: TTTCCGTGTTTTGTTTCTCATCG (forward) /TGTTGTTCTTGTTGTTGTTGTCA (reverse) at 3R 7,215,009 and TGATGTTGATGAGCGCACAG/AAATGCTGTCACACGCTTTG at 3R 6,883,792. We also designed three pairs flanking indels inside the distortion region: CCAGGTGGGTACTCAATAGATTT/CTGTTGGAAATGGAGGTGAGAA at 3R 22,400,780, GGCTCTGGGCCATGTCAATA/GTGCGTGTTGGCCTGTTAAT at 3R 23,929,097, and CTGGGGAGTAGCACGTTTCC/GATGTGGATGTGGCTGTGGA at 24,917,176.

## Acknowledgments

We thank Dr. P. Satyaki for the HeT-A probe and Dr. Dean Castillo, Dr. Anne-Marie Dion-Côté, Dr. Alla Kalmykova, Dr. Sarah Sander and the anonymous reviewers for helpful comments. Supported by NIH R01 GM074737 to D.A.B. and R01 AI064590 to A.G.C.

---

**Supplementary Figure 1.**
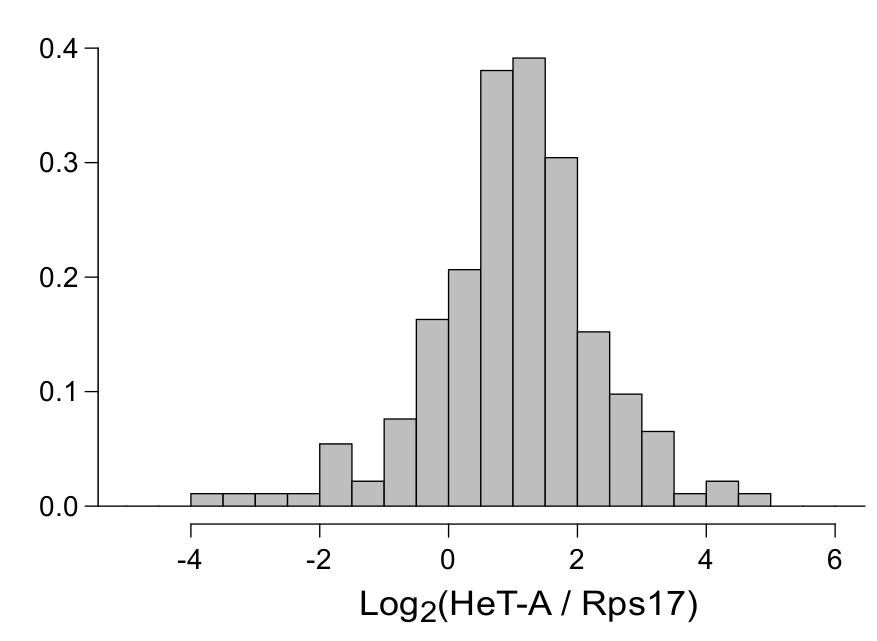
Histogram of HeT-A quantities in log scale based on data from Fig. 1A.

**Supplementary Figure 2.**
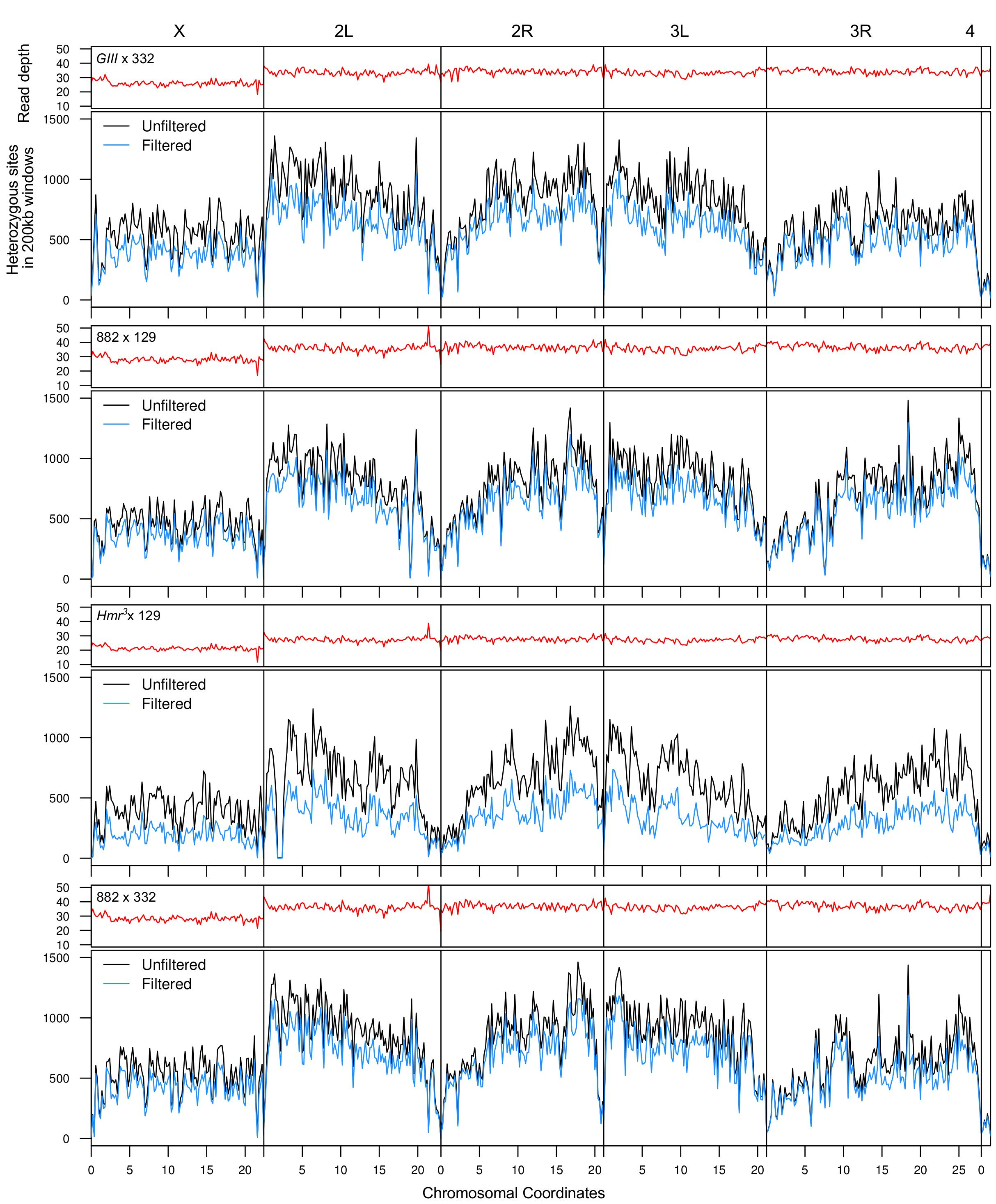
Read depth and SNP density. For each of the four crosses, the read depth across the genome is plotted in the top panel, and the number of heterozygous SNPs before and after filtering by read depth in each 200kb window is plotted in the lower panel. The read-depth filter removes SNPs with read depths below 20 (X-linked) or 25 (autosomal), or above twice the average read depth.

**Supplementary Figure 3.**
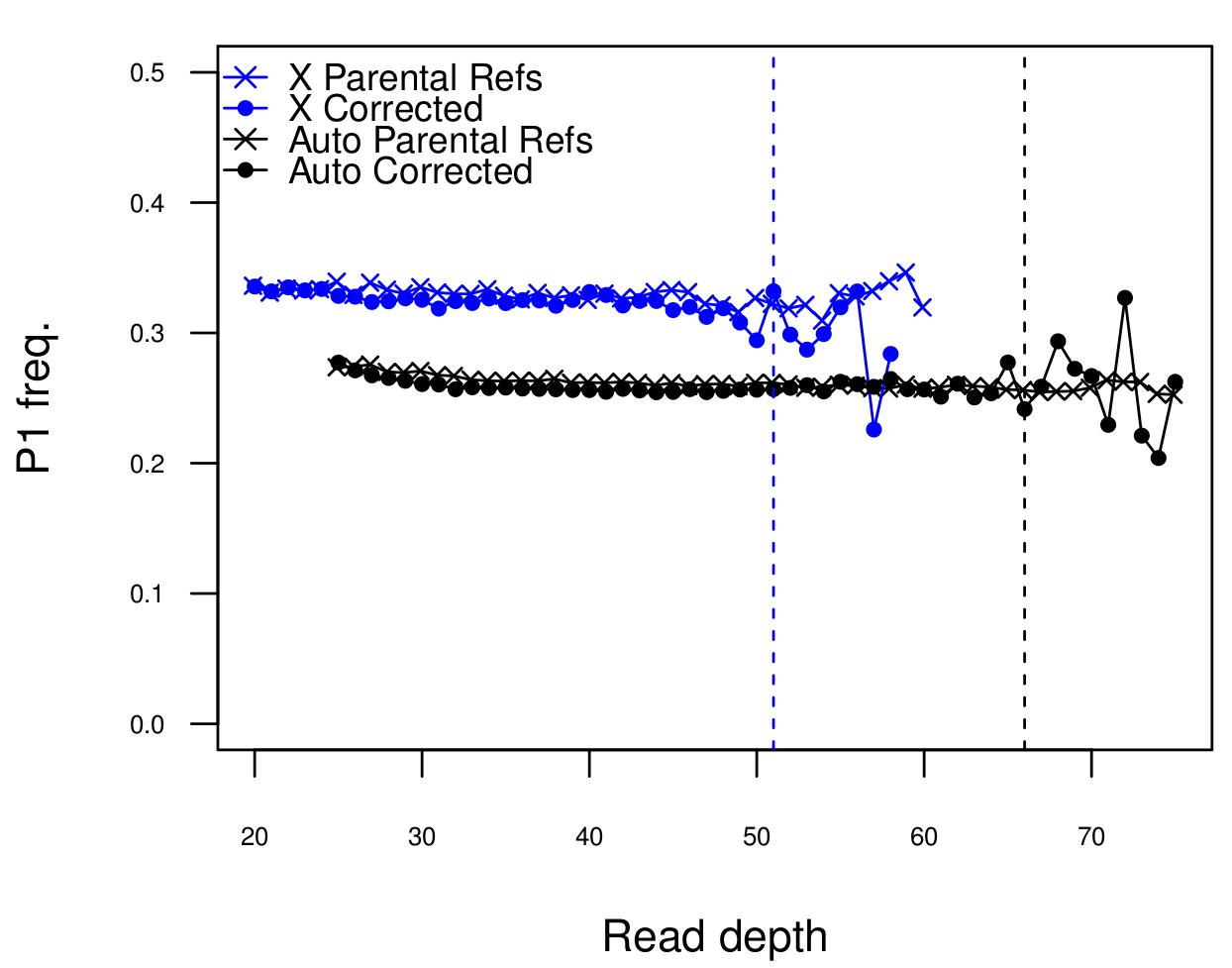
Reference allele bias correction. The allele frequencies after reference allele bias correction (replotted from Figure 2C) are plotted along with allele frequencies generated from alignments to both parental genomes, an alternative strategy to minimize bias. To do this, we used GATK’s Alternate Reference Maker to modify the reference genome with SNPs found in DGRP-882 or DGRP-129, generating the two parental genomes to which the DGRP-882 x DGRP-129 cross was aligned. After SNP calling with GATK, the P1 and P2 counts at each heterozygous site are simply the reference counts from the alignment to DGRP-882 and DGRP-129, respectively. The dashed vertical lines represent the upper-bound read depth cut off (twice the average read depth).

**Supplementary Figure 4.**
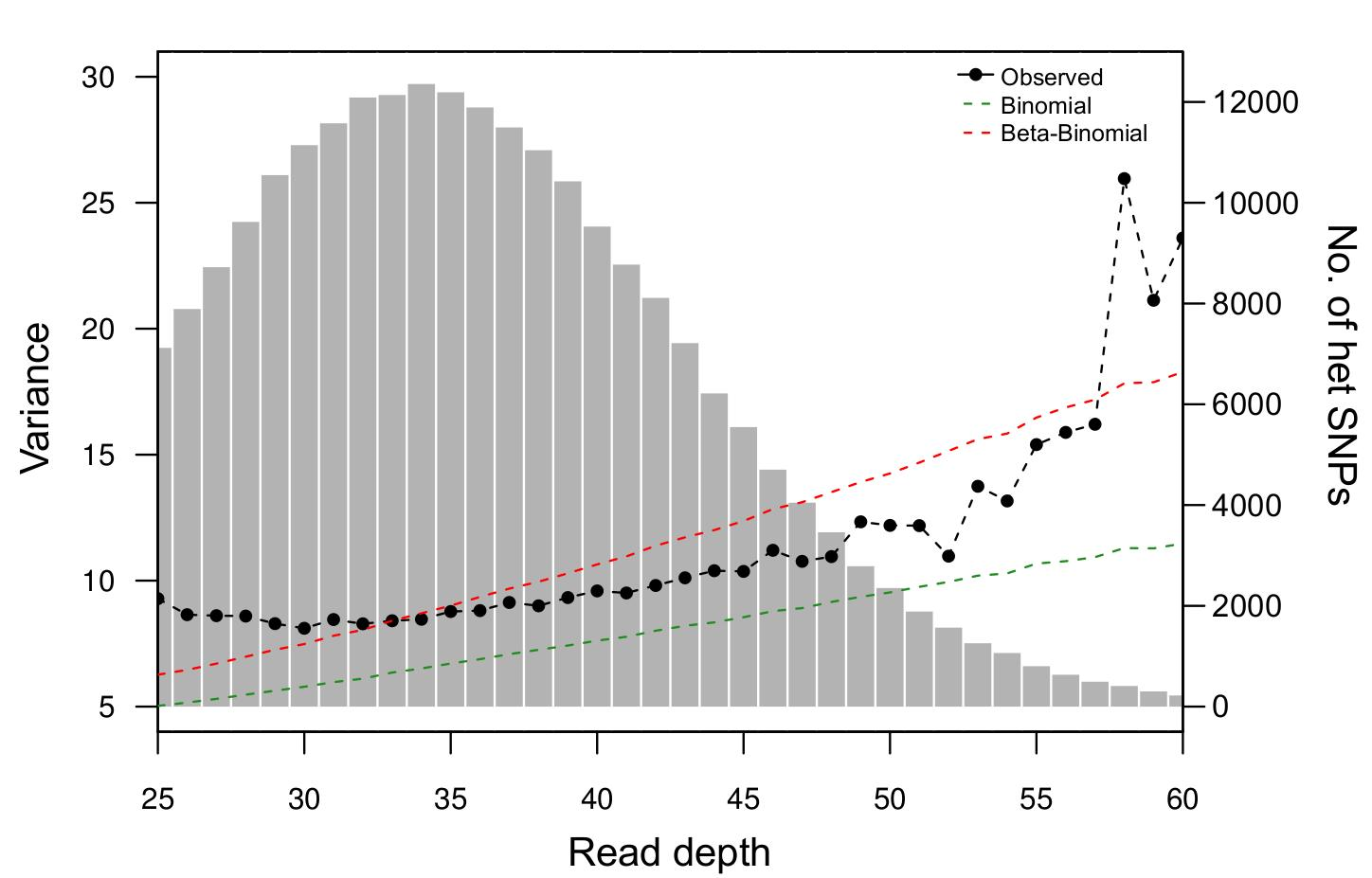
Estimating and modeling variance in allele frequency. The observed variance of P1 counts at different read depth bins is plotted with black connected dots. The expected variance under Binomial (green) and Beta-Binomial (red) sampling is plotted in dotted lines. The histogram of the read depth distribution of heterozygous SNPs is plotted with gray shading in the background.

**Supplementary Figure 5.**
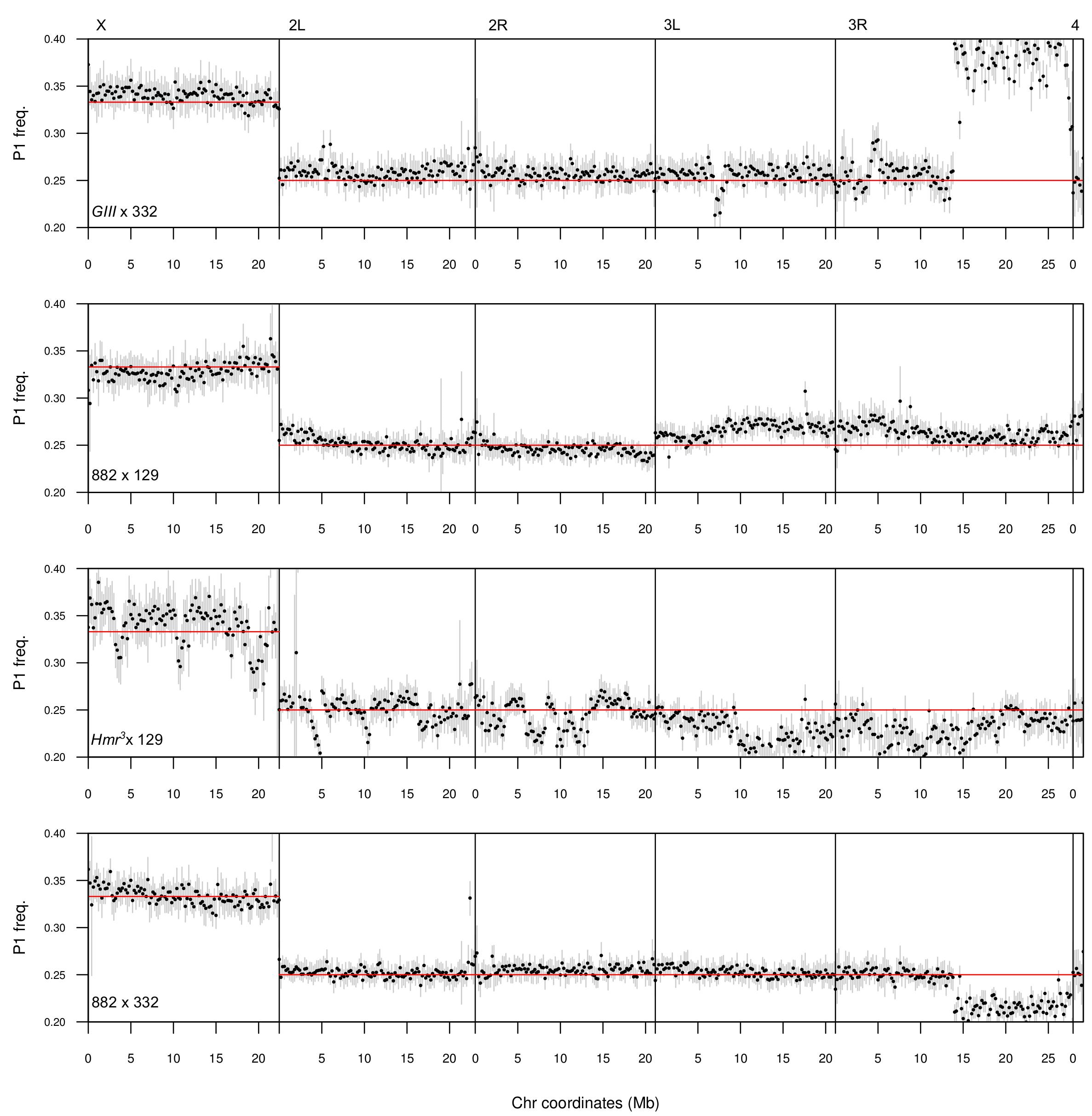
Allele frequency across chromosomes after filtering out SNPs within 100bps. Because heterozygous sites that are less than 100bps apart cannot be independently sampled given 100bp reads, sites used to generate Figure 3 were further filtered to remove neighboring SNPs within 100bps. When multiple heterozygous SNP sites were found within 100bp, only one site was kept at random.

**Supplementary Figure 6.**
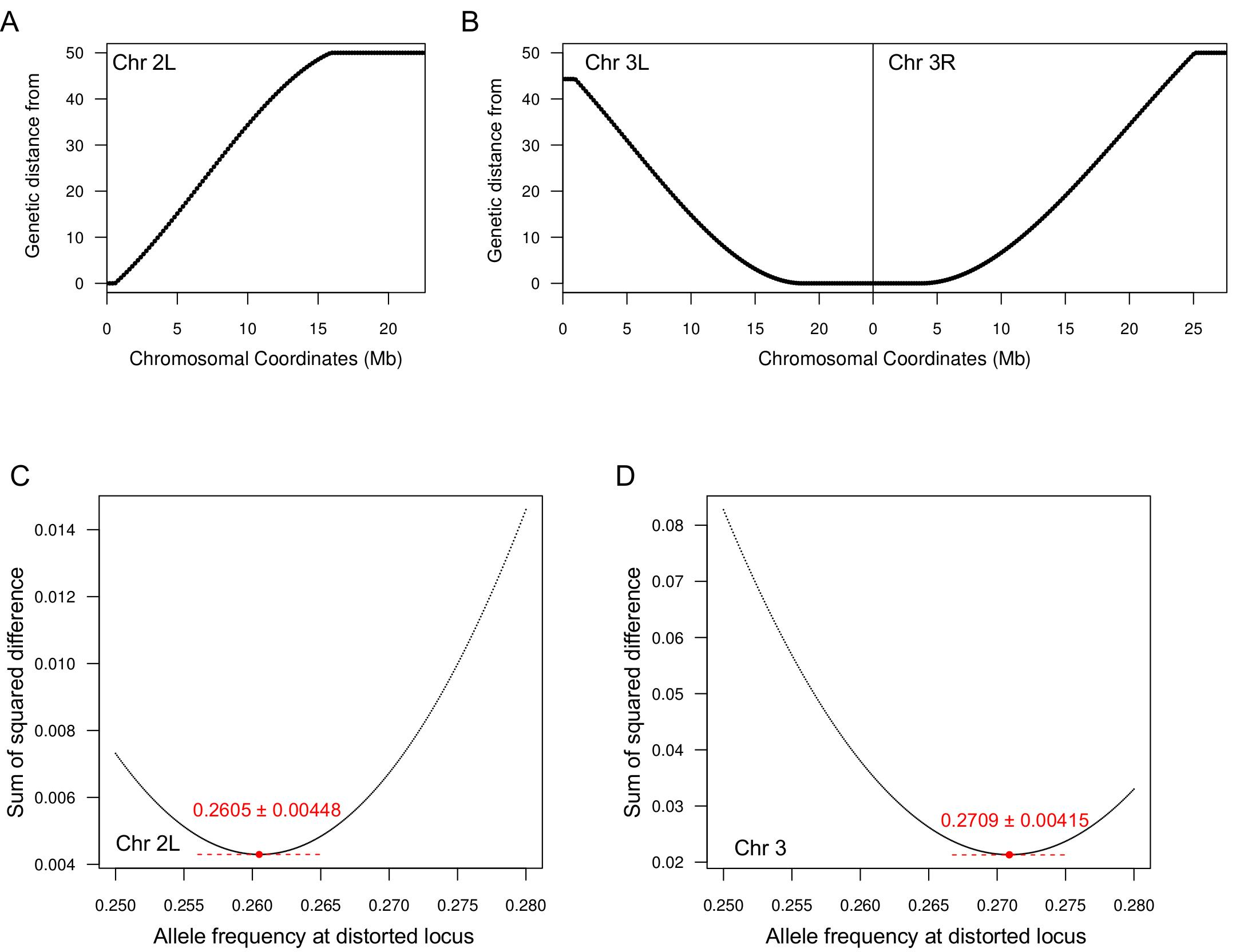
Distortion estimates based on best fit to recombination decay. Genetic distance from the telomere of Chr. 2L (A) and centromere of Chr. 3 (B) is plotted. To estimate the allele frequency at the two putative distortion loci from the DGRP-882 x DGRP-129 cross in Figure 5, we determined the fit of the decay curve with distorted P1 allele frequencies increasing at 0.0001 intervals, starting at 0.25 (i.e. no distortion). The fit as determined by the sum of squared difference between the observed allele frequency at each window and the curve is plotted for each distortion allele frequency. The least square value is approximated by the lowest point (red) in the graphs. The dotted red lines demarcate the 95% confidence intervals for the least square estimates which are also noted in red above of the points. For the locus at the telomere of Chr. 2L (C), the fit was based only on chromosome arm 2L because arm 2R is unlinked from the telomere. For the locus at the centromere of Chr. 3 (D), the fit was summed over both 3L and 3R.

**Supplementary Figure 7.**
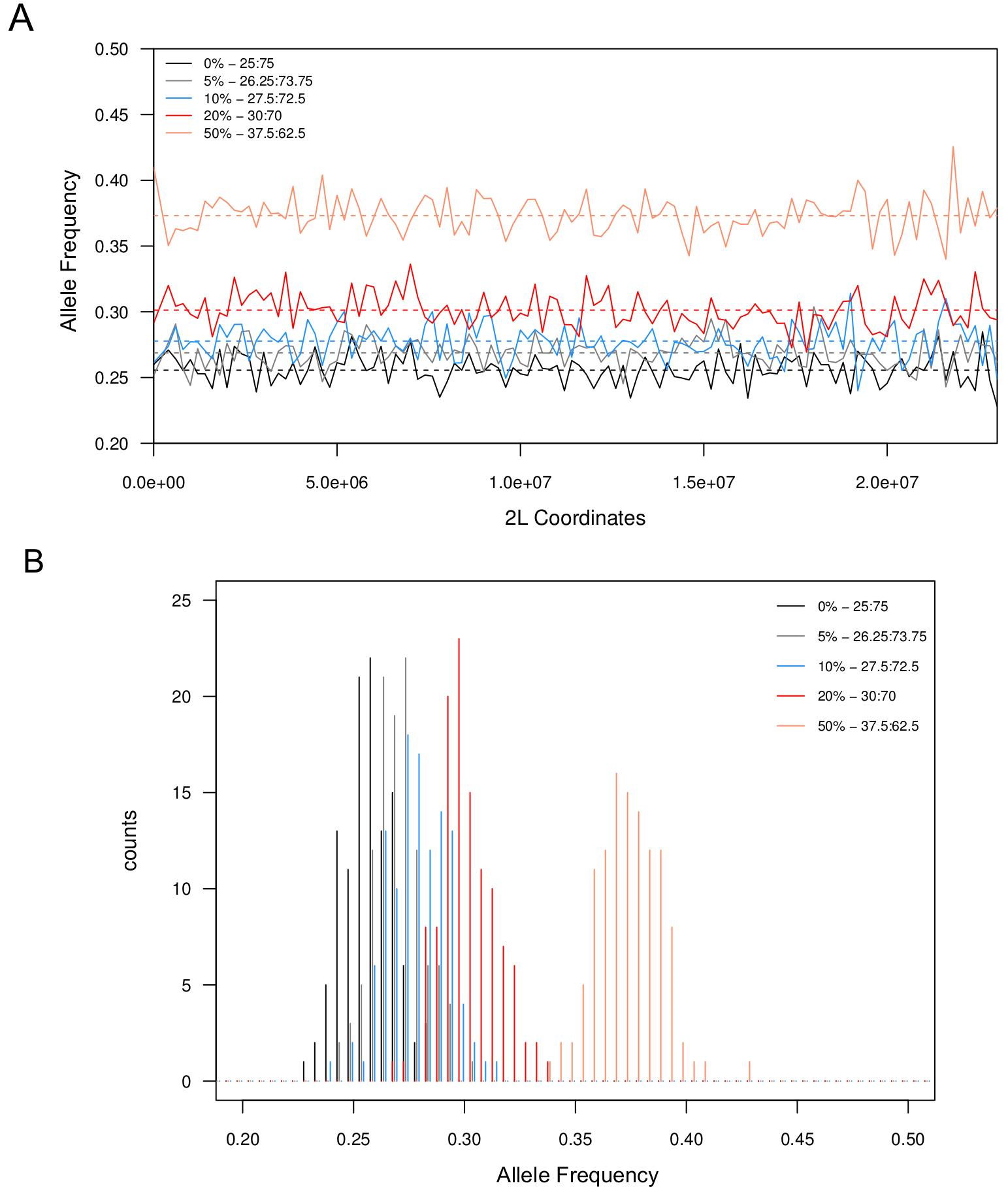
Simulation of crosses with meiotic drivers of different strengths. We simulated the results of pooled sequencing by generating and concatenating simulated reads from chromosome 2L of DGRP-882 and DGRP-129 at different ratios. The reads were then mapped/processed as per our pipeline. A) With drivers of different strengths (different colors as indicated by legend), P1 allele frequencies in 200kb windows are plotted across the chromosome for one simulation each. Dotted lines indicate the average across the chromosome for each driver strength. B) Distributions of allele frequencies for all 200kb windows plotted in A) above for different meiotic driver strengths. All pair-wise comparisons of the distributions are statistically significant (Wilcoxon Ranked Test).

